# New waves: Rhythmic electrical field stimulation systematically alters spontaneous slow dynamics across mouse neocortex

**DOI:** 10.1101/171926

**Authors:** Anastasia Greenberg, Javad Karimi Abadchi, Clayton T. Dickson, Majid H. Mohajerani

**Affiliations:** Neuroscience and Mental Health Institute, University of Alberta, Edmonton, AB, Canada, T6G 2E1; Canadian Centre for Behavioural Neuroscience, University of Lethbridge, Lethbridge, AB, Canada, T1K 3M4; Department of Psychology, University of Alberta, Edmonton, AB, Canada, T6G 2H7; Department of Physiology, University of Alberta, Edmonton, AB, Canada, T6G 2E9

## Abstract

The signature rhythm of slow-wave forebrain activity is the large amplitude, slow oscillation (SO: ~1 Hz) made up of alternating synchronous periods of depolarizing and hyperpolarizing states at the single cell and network levels. On each wave, the SO originates at a unique location and propagates across the neocortex. Attempts to manipulate SO activity using electrical fields have been shown to entrain cortical networks and enhance memory performance. However, neural activity during this manipulation has remained elusive due to methodological issues in typical electrical recordings. Here we use voltage-sensitive dye (VSD) imaging in a bilateral cortical preparation of urethane-anesthetized mice to track SO cortical activity and its modulation by sinusoidal electrical field stimulation applied to frontal regions. We show that under spontaneous conditions, the SO propagates in two main opposing directional patterns along an anterior lateral – posterior medial axis. Rhythmic field stimulation alters spontaneous propagation to reflect activity that repeats cycle after cycle with distributed and varied anterior initiation zones and a consistent termination zone in the posterior somatosensory cortex. Our results show that slow electrical field stimulation stereotypes ongoing slow cortical dynamics during sleep-like states.

**Author Contributions:** A.G., J.K.A., M.H.M. and C.T.D. designed the study. A.G. and J.K.A. performed the experiments. A.G. analyzed the data. A.G. wrote the manuscript, which all authors commented on and edited. C.T.D. and M.H.M. supervised the study.

---

Synchronized brain rhythms during distinct behavioral states such as sleep and wakefulness are increasingly recognized as paramount for physiological processing ^1, 2, 3, 4, 5^ ^6^. Within sleep and anesthesia, cortical activity is marked by the large amplitude rhythmic slow oscillation (SO; ~ 1 Hz) at the network level which is generated by toggling between depolarized (ON, UP) and hyperpolarized (OFF, DOWN) states at the cellular level^7, 8, 9^. This rhythm has been implicated in hippocampal-dependent memory consolidation in both human ^10, 11, 12^ and animal studies ^13, 14, 15^. The SO has several properties that make it suitable for such a role including its grouping of faster local gamma rhythms ^16, 17^, thalamocortical spindles ^18, 19^ and hippocampal memory replay events – sharp-wave ripples^20, 21, 22^ – into its ON state, allowing for cross-network interactions. On any given wave, the SO originates at a specific cortical focus and travels across the cortex ^23, 24^ to the hippocampus ^25, 26, 27^. This activity generally propagates in an anterior-to-posterior direction from many possible initiation zones with many complex pattern variations ^23, 24, 28, 29, 30^.

Attempts to enhance the SO using slow alternating electrical stimulation applied to frontal cortical regions have resulted in improved hippocampal-dependent memory performance in both humans ^10, 31^ and rats ^32^. However, the mechanism by which such stimulation affects SO dynamics in favour of improved memory performance is not well understood. It has been demonstrated that rhythmic field stimulation boosts subsequent SO power, promotes hippocampal ripples, and entrains both cortical and hippocampal activity ^10, 33, 34, 35^. Simple electrode array experiments have shown that field stimulation biases and stereotypes SO propagation ^33^. However, a more nuanced understanding of how stimulation influences large-scale cortical dynamics is forthcoming.

Using large-scale voltage-sensitive dye (VSD) optical imaging of a large bilateral cortical window during slow-wave conditions, we examined spontaneous and electrical-field modulated dynamics. We show that the typical pattern of spontaneous activity and its propagation along an anterior-posterior axis is profoundly altered and stereotyped by the influence of entraining rhythmic electrical fields. This is an important demonstration of the power of mesoscale population imaging to identify activity-dependent alterations of spatially broad neural networks during field applications that would interfere with typical electrophysiological methods; and brings us closer to understanding how these manipulations may produce their complex influences on behavior.

## Results

### Large scale spontaneous slow-wave cortical activity patterns show broad and characteristic motifs

We employed voltage-sensitive dye (VSD) imaging of a large bilateral cortical window in urethane-anesthetized mice to assess the effects of frontally-applied sinusoidal electrical field stimulation on subthreshold slow-wave dynamics ^36^. We imaged cortical dynamics using a fast CCD camera with high spatial (67 µm per pixel) and temporal (5 – 10 ms) resolution (Figure 1A) following dye application to the surface of the brain (see Methods) ^37, 38^ ^39^.

**Figure 1.**
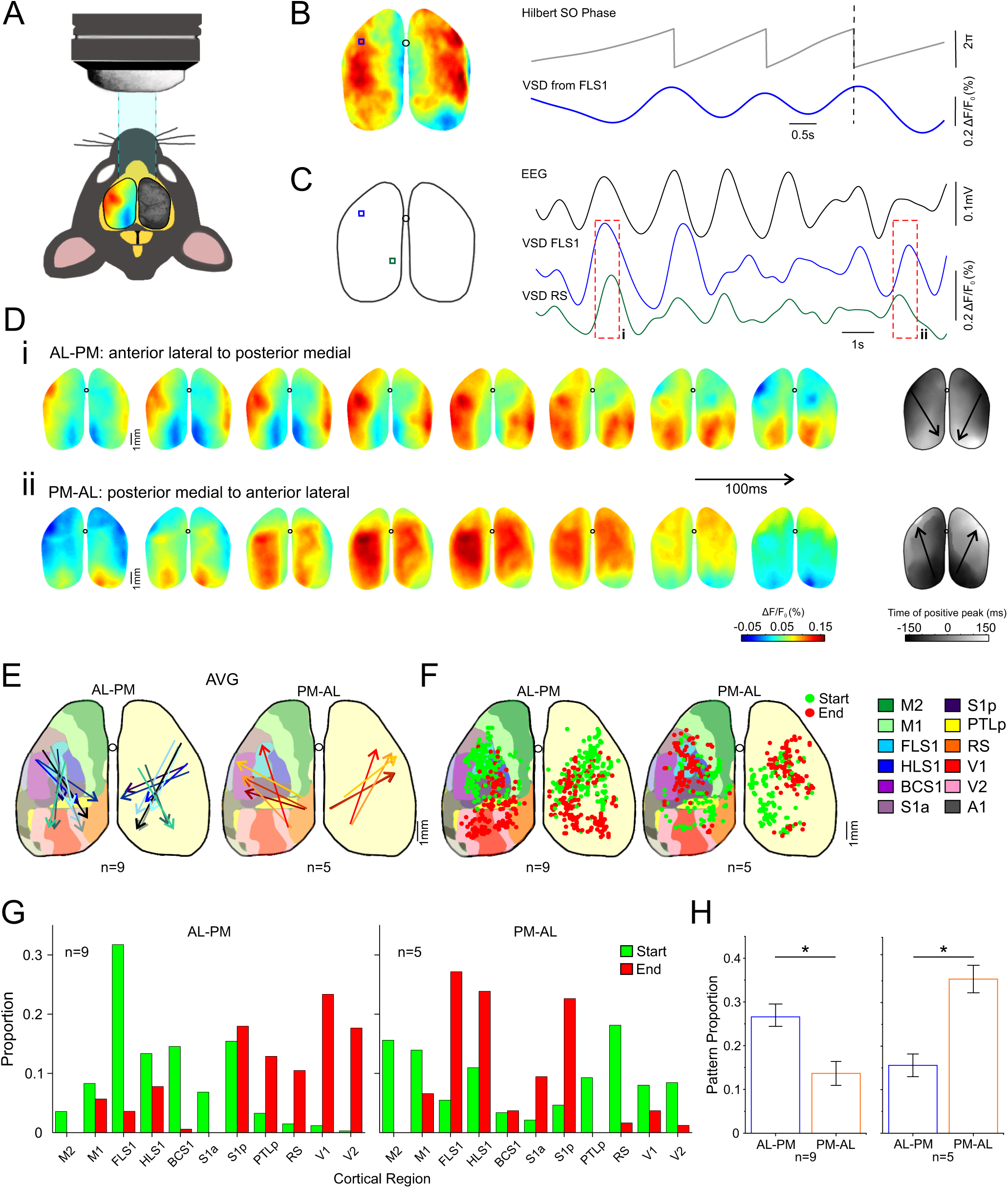
The spontaneous slow oscillation propagates in two major opposing directional patterns along an anterior-lateral – posterior-medial axis. **A.** Experimental set-up with mouse in aerial view. Left hemisphere shows VSD signal and right hemisphere shows camera image of exposed cortex. **B.** Left: VSD image frame taken at dotted line on right. Blue box indicates ROI at FLSI. Right: VSD trace from FLS1 and corresponding Hilbert phase angle. **C.** Traces on right for EEG, and VSD ROI at FLS1 and RS (retrolental) shown on brain map on left. **D.** Colored images show VSD maps filtered for the slow oscillation range (0.3 – 1.5 Hz) taken at time points during a single cycle for i and ii shown by red boxes in C. In i the given cycle showed a propagation pattern with activity (positive; warm colors) starting from anterior-lateral regions and ending in posterior-medial regions. In ii the opposite pattern is apparent with activity starting in posterior-medial regions and ending in anterior-lateral regions. Grey images on the right show time lags of positive peaks in VSD signals for each pixel for the given propagation cycle. Positive peaks were identified using Hilbert phase and a lag of zero indicates positive peak at the ROI of HLS1 (top brain inset). Arrows show average gradient direction from the earliest to latest time lags. **E.** Average trajectory arrows as in A across cycles for the single major pattern for every experiment plotted on standardized cortical map. Left map shows arrows for 9 experiments in which the AL-PM pattern was the major pattern, and right map shows the remaining 5 experiments in which the PM-AL pattern was the major pattern. **F.** Start and end points of activity propagation plotted on standardized cortical maps for every cycle of the major pattern in every experiment separated by whether the major pattern was an AL-PM or PM-AL pattern (left and right maps respectively). Legend indicates cortical areas represented in color on standardized maps. **G.** Proportion of start and end points from C represented in each cortical region. **H.** Proportion of all patterns that the major and second major patterns represent across experiments separated by whether the major pattern was an AL-PM or PM-AL pattern.

To describe the influence of field stimulation on SO propagation dynamics, it was necessary to systematically describe spontaneous activity conditions in our preparation. We first mapped the cortical surface being recorded via multimodal sensory-stimulation in order to functionally delineate regions (Supplementary Figure 1). Our spontaneous VSD recordings across regions showed characteristic bilaterally synchronous SO activity (~1 Hz) that corresponded well to simultaneously recorded EEG (Figure 1B,C; Supplementary Figure 2).

We confirmed that both our spontaneous and stimulation-evoked recordings reflected neural activity by showing that they were abolished during conditions of sodium channel blockade (lidocaine), glutamatergic antagonism (CNQX+MK-801), as well as during post-mortem conditions (Supplementary Figures 1,2).

We mapped propagation patterns using measures of instantaneous SO phase on a cycle-by-cycle basis in order to track the peak ON state activity across all imaged pixels and clustered these patterns into groups using a hierarchical clustering algorithm (Figure 1B; see Methods). During baseline conditions, two major patterns emerged across all experiments out of an average of 14±1 patterns detected (Figure 1). The most common patterns, in all animals, traveled bidirectionally along an anterior to posterior axis. The most common pattern across animals began in an anterior-lateral position and propagated in a quasi-linear fashion to posterior-medial regions. We called this anterior lateral-to-posterior medial (AL-PM) propagation. The second most common pattern was the mirrored opposite (posterior to anterior) with initiation in posterior-medial regions and propagation to anterior-lateral positions. We called this posterior medial-to-anterior lateral (PM-AL) propagation (Figure 1; Supplementary Movies 1,2). Although the AL-PM pattern was the most common overall, in five of nine animals the PM-AL pattern that was most common, with the AL-PM pattern a close second. These two patterns together represented 44±2% of all detected patterns across all experiments (Figure 1H).

We also assessed the initiation and termination points of propagation for every SO cycle for each of the two major patterns across experiments by detecting the center of mass of the pixels with the earliest and latest Hilbert supra-threshold positive peak detections (see Methods). As shown in Figure 1 (panels F&G), the spatial distribution of both initiation and termination points was variable. The AL-PM pattern showed a high proportion of initiation points in somatosensory areas (81% in S1 generally and especially in FLS1: 32%), with termination points mainly in posterior regions (64% across V1, V2, pTLP, and RS) (Figure 1G – left). For the PMAL pattern on the other hand, the initiation points were more broadly distributed across posterior medial areas including retrosplenial, posterior parietal, primary and secondary visual cortices with the majority located outside of somatosensory areas (73%) while end points were mainly within the somatosensory region (S1, 87%) (Figure 1G – right). Pattern likelihood was not correlated with the amplitude of the VSD SO on individual cycles (Supplementary Figure 3).

We noted several unique propagation patterns under spontaneous conditions that were less common but were nevertheless present across experiments: patterns with multiple initiation points, multiple termination points, initiation and termination points located in posterior regions of the brain, and asymmetrical patterns with initiation/termination points appearing with a time lag across hemispheres (Supplementary Figure 4).

### Field stimulation alters and entrains VSD slow-wave activity

Application of slow (1.67 Hz) alternating electrical fields suppressed VSD SO power and RMS in comparison to spontaneous recordings (0.5 – 0.8 Hz; Power: baseline = 0.005±0.001, STIM = 0.002±0.001; p = 0.03; RMS: baseline = 0.039±0.004, STIM = 0.018±0.002; p = 0.002; Figure 2B-D). In contrast, both power and RMS at the frequency of stimulation (1.67 Hz) increased (although non-significantly: Power: baseline = 0.001±0.00, STIM = 0.002±0.001; p = 0.19; RMS: baseline = 0.015±0.001, STIM = 0.018±0.002; p = 0.054; Figure 2B,C,E) in every experiment, suggesting entrainment. Interestingly, this suppression of spontaneous SO continued following termination of the stimulation and appeared to be correlated with the intensity of the applied current (Supplementary Figure 5,6).

**Figure 2.**
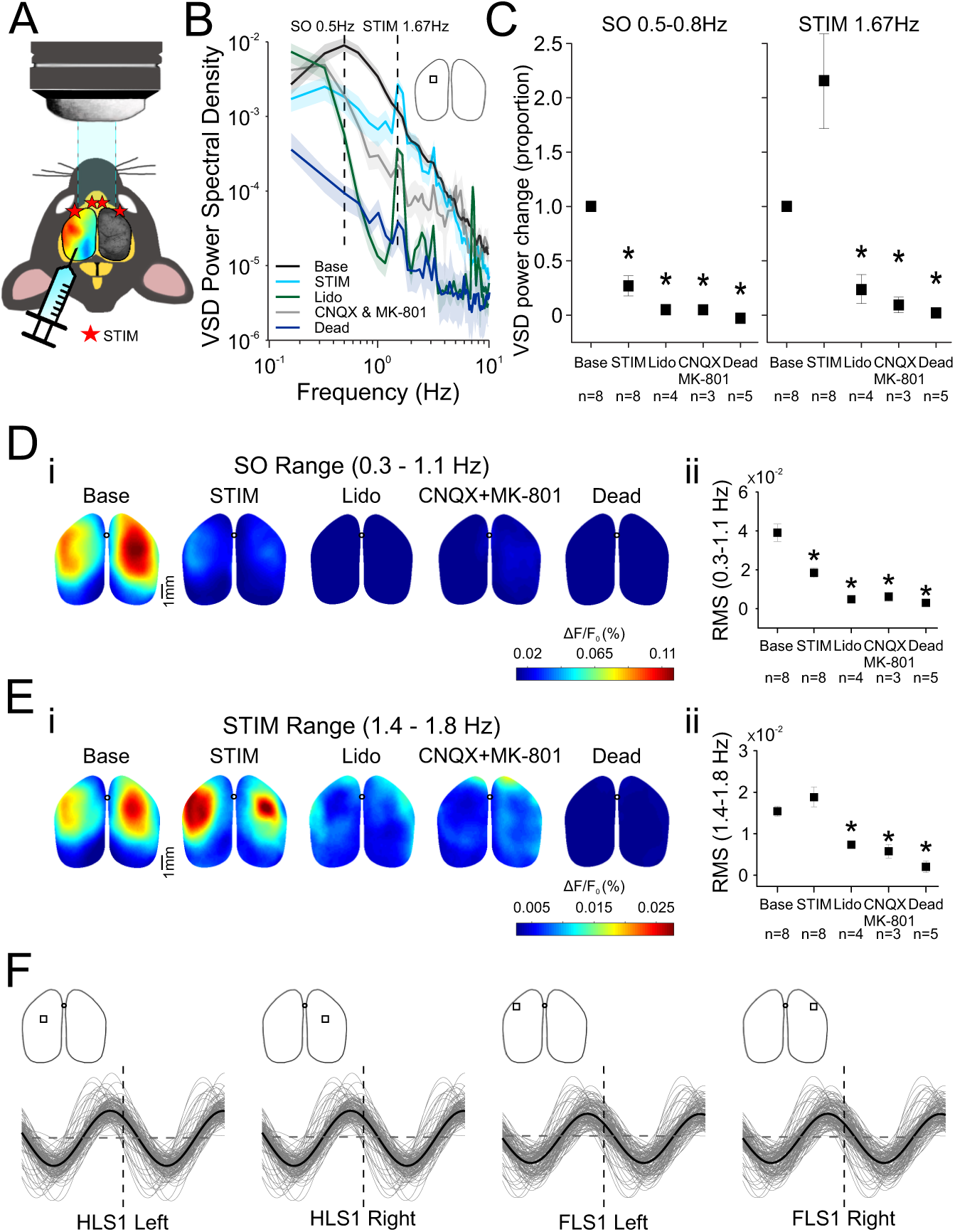
Figure 2 Sinusoidal field stimulation alters slow-wave EEG and VSD spectral properties. **A.** Experimental set-up showing locations of stimulation (stars). **B.** VSD power spectra during baseline as well as during the application of field stimulation under 4 conditions: unsuppressed (STIM), lidocaine, CNQX+MK-801 and following death. Brain inset shows location of ROI used for power calculation. Dotted lines indicate peak baseline slow oscillation frequency (0.5 Hz) and the sinusoidal stimulation frequency (1.67 Hz). Note spectral peaks at the exact frequency of stimulation for all stimulation conditions with lower power in the drug and death conditions compared to the STIM condition. Stimulation frequency peaks are magnitudes of order weaker power compared to the unsuppressed STIM condition. **C.** Proportion of VSD power change from baseline across experiments for the peak slow oscillation frequency (0.5 – 0.8 Hz depending on the experiment) and for the stimulation frequency (1.67 Hz) for all conditions. Note weaker slow oscillation power for all stimulation conditions and a non-significant boost in the stimulation frequency power specific to the STIM condition (without activity suppression). **D.** i. RMS VSD maps computed for the slow oscillation range (0.3 – 1.1 Hz) on three-minute segments for all conditions in the same experiment as in B. **ii.** RMS for the slow oscillation range across experiments. **E. i.** RMS VSD maps computed for a frequency range (1.4-1.8 Hz) that includes the stimulation frequency (1.67 Hz) for the same experiment as in B and C. **ii.** RMS for the stimulation range across experiments. Note weaker slow oscillation RMS during sine-wave stimulation (STIM) and non-significant boost in RMS during for the stimulation range. Also note that the artifact component RMS that appears for the stimulation range during activity suppression conditions is much weaker and spatially distinct when compared to the unsuppressed STIM condition. **F.** Single VSD traces (gray) and average trace (black) taken from ROIs indicated by brain insets triggered the peak of the stimulation sine-wave during stimulation. All traces are from the same experiment. Vertical dotted line indicates trigger time point at the positive peak of the stimulation wave. Horizontal dotted lines indicate 95^th^ percentile confidence intervals based on triggered-averages of randomized time points.

To confirm that VSD signals were not contaminated by the applied electrical fields themselves, we compared VSD signals before and after application of lidocaine, glutamate receptor antagonism, as well as pre- and post-mortem. Under these conditions, VSD power and RMS values were magnitudes of order lower than for spontaneous condition for both the SO range (VSD: baseline = 0.005±0.001, lidocaine = 5.4E-5±1.3E-5, p = 0.016; CNQX+MK-801 = 9.5E-5±4.6E-5, p = 0.035; Dead = 5.6E-5±1.8E-5, p = 0.008; RMS: baseline = 0.039±0.004, lidocaine = 0.005±0.001, p = 0.004; CNQX+MK-801 = 0.006±0.002, p = 0.003; dead = 0.003±0.002, p < 0.001; Figure 2C,D) and at the stimulation frequency (VSD: baseline = 0.001±9.8E-5, lidocaine = 5.1E-4±2.2E-4, p = 0.003; CNQX+MK-801 = 1.3E-5±7.1E-7, p = 0.004; dead = 7.0E-5±4.5E-5, p < 0.001; RMS: baseline = 0.015±0.001, lidocaine = 0.007±0.001, p = 0.004; CNQX+MK-801 = 0.005±0.001, p = 0.004; dead = 0.002±0.001, p < 0.001; Figure 2C,E) demonstrating that any artefactual component is highly unlikely to affect our signal of interest. Importantly, this demonstrates that global cortical activity dynamics can be successfully resolved during the application of electrical fields, a result that has defied other classic electrophysiological methods.

Given the tendency for VSD spectral power to be boosted at the frequency of stimulation, we assessed the level of VSD activity entrainment to the phase of stimulation. There was strong entrainment of VSD activity to the stimulus sine wave during stimulation for all locations selected (bilaterally: HLS1, FLS1, V1). The degree of phase entrainment was based on a comparison to randomized trigger points (p < 0.001; Figure 2F, Supplementary Figure 7). The level of entrainment to the sine wave was similar in magnitude to the baseline VSD entrainment to the spontaneous SO (Supplementary Figure 5). When applying stimulation during activity-suppressing conditions (lidocaine, CNQX+MK-801, post-mortem), triggered averages had negligible (and significantly lower) sine-wave entrainment (Supplementary Figure 7).

### Field stimulation alters and stereotypes slow oscillation propagation dynamics

To assess any alterations of slow-wave dynamics via imposed electrical fields, we tracked propagation patterns cycle-by-cycle during stimulation conditions by focusing on the stimulation frequency (1.67 Hz; VSD bandpass filtered between 1.1 and 1.9 Hz). Before stimulation we observed the AL-PM and the PM-AL propagation patterns for spontaneous SO as described above (c.f. Figure 1). We also confirmed that propagation during spontaneous conditions at the stimulation frequency resembled that of the ~1Hz SO. During stimulation, propagation was altered and the major propagation pattern detected did not overlap with any existing patterns during baseline at either frequency (Figure 3; Supplementary Movie 3). On any given cycle, activity could initiate at multiple locations but consistently terminated within the somatosensory region (S1; 65%), with the largest proportion in the posterior somatosensory area (S1p, 32%) (Figure 3). This region borders the posterior parietal associational area (PTLp) and includes the somatosensory trunk area (TrS1) and an unassigned area (UnS1) (Figure 3B,C – S1p, dark purple). Overall, the major pattern during stimulation appeared to be propagating in an anterior-to-posterior direction, in line with our previous work ^33^. To confirm this, when we constrained propagation to three points along an anterior-posterior axis in M1, we show that stimulation tends to bias propagation in an anterior-posterior direction (Supplementary Figure 8)^33^.

**Figure 3.**
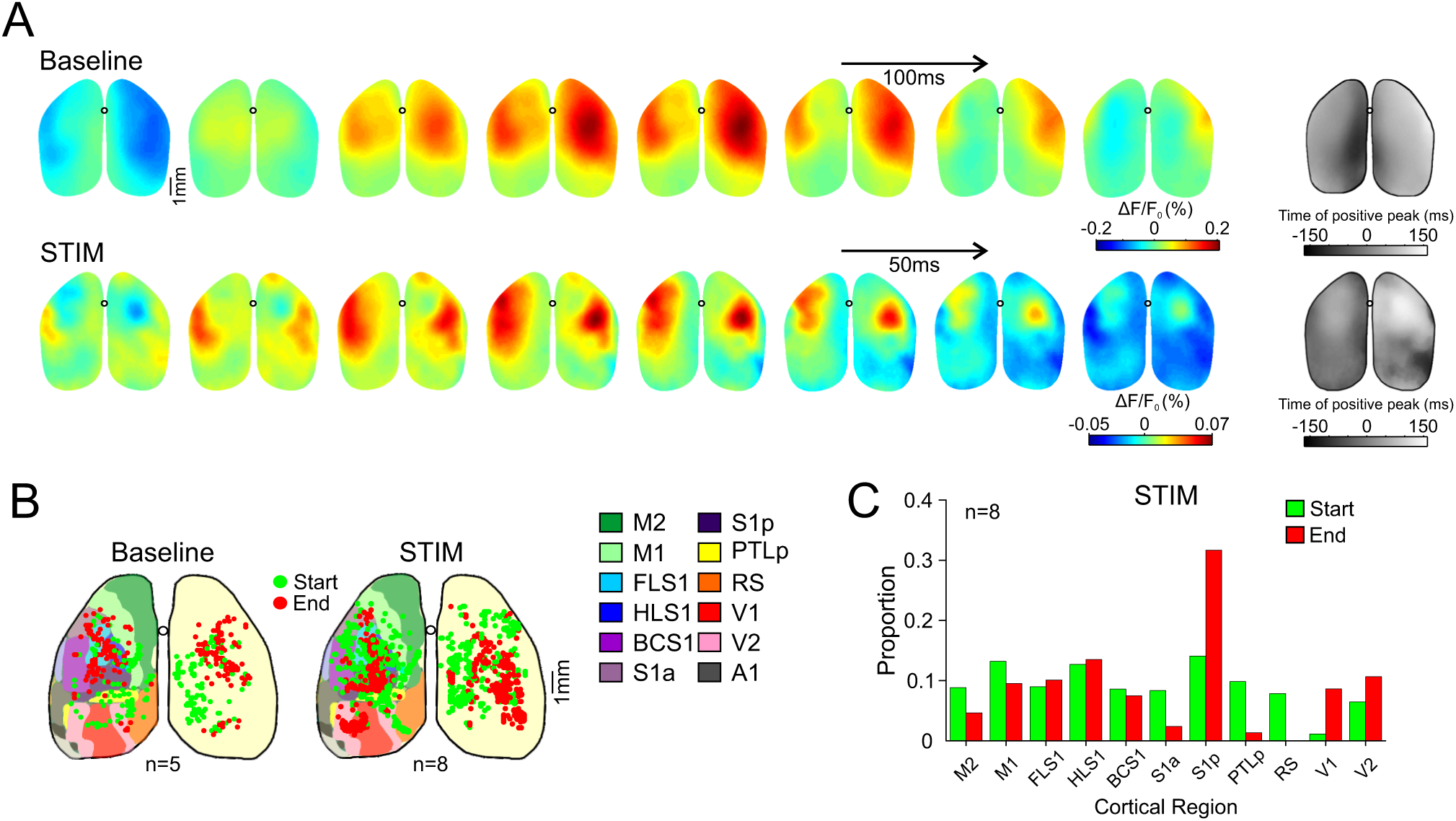
Sinusoidal field stimulation alters slow oscillation propagation dynamics. **A.** Colored images show VSD maps filtered for the slow oscillation range (0.3 – 1.5 Hz) for baseline or for the stimulation range (1.5 – 2.0 Hz) for the stimulation conditions taken at 8 time points during a single cycle. For each condition, the cycle depicted is taken from the major pattern group detected. The baseline cycle shows a PM-AL propagation pattern which was the major pattern in this experiment. During stimulation, the major propagation pattern appears altered. Grey images on the right show time lags of positive peaks in VSD signals for each pixel for the given propagation cycle. Positive peaks were identified using Hilbert phase and a lag of zero indicates positive peak at the ROI of HLS1 for baseline and peak of stimulation sine wave for all other conditions. **B.** Start and end points of activity propagation plotted on standardized cortical maps for every cycle of the major pattern for both conditions in every experiment. The baseline map is the same as shown in Figure 1 for the PM-AL pattern and chosen to reappear here as this was the major pattern for the experiment shown in A. Note altered distribution of start and end points for the STIM condition that does not resemble either of the two major baseline patterns (c.f. Figure 1). **C.** Proportion of start and end points from B represented in each cortical region for the stimulation condition.

To confirm that our results were not as a result of artefactual imposition of alternating electrical fields, we also assessed slow-wave dynamics during conditions of suppressed activity (lidocaine, CNQX+MK-801 and post-mortem). As before, during these conditions, VSD activity was enormously weaker than during unsuppressed stimulation and had an exceedingly different spatial profile with initiation and termination points distributed around the edges of the craniotomy window, reflecting the pure artefactual component of stimulation (Supplementary Figure 9).

Field stimulation not only altered SO propagation patterns but also stereotyped its activity creating a less dynamic oscillation. During baseline conditions, our clustering algorithm detected 14±1 propagation patterns which decreased significantly during stimulation to 7.6±1 patterns (p < 0.001; Figure 4A – left). Likewise, the proportion of all patterns represented by the single major pattern doubled during stimulation (52±5%) compared to baseline (25±2%) (p < 0.001; Figure 4A – right). There was no change during stimulation in the propagation index (change in area of activation per frame; p = 0.34) or velocity of propagation (p = 0.44) (Figure 4B). Importantly, stimulation applied during lidocaine, glutamate receptor antagonism and postmortem showed significantly lower propagation index values (lidocaine: p = 0.02; CNQX+MK-801, p = 0.043; dead: p = 0.002) compared to baseline (Figure 4B). This demonstrates that there was no true propagation of activity during neural suppression.

**Figure 4.**
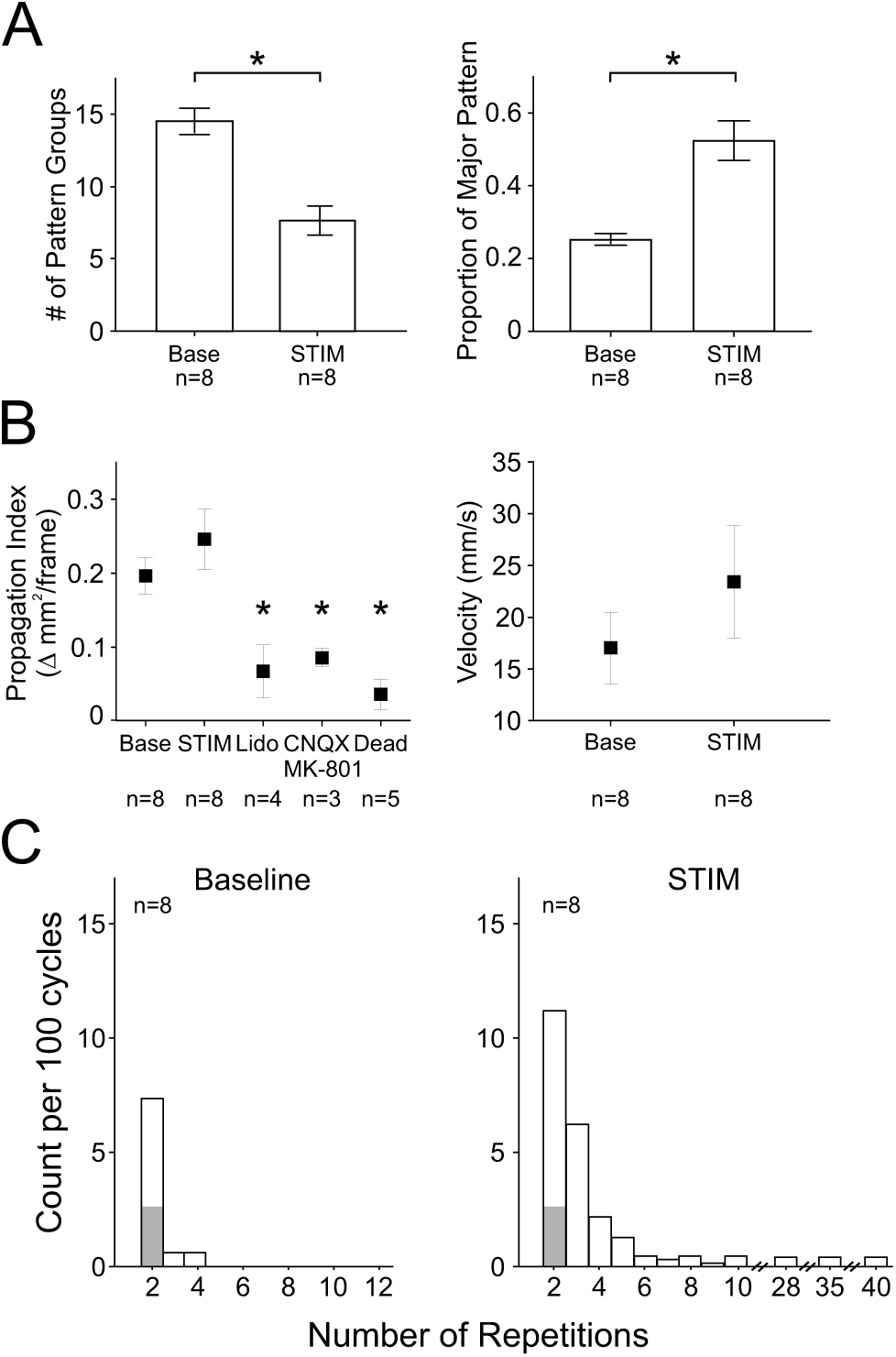
Sinusoidal field stimulation stereotypes propagation patterns. **A.** Left: The number of pattern groups identified with the hierarchical clustering algorithm across experiments for the baseline and stimulation conditions. Right: The proportion of all patterns that the major pattern represents across experiments for the baseline and stimulation conditions. **B.** Left: propagation index as a change in area of activation (number of pixels >2SD of VSD signal) per frame across experiments for baseline, stimulation and the three activity suppressing conditions. Note significantly lower propagation index values in the activity suppressing conditions suggesting that these conditions show static artifact which does not propagate over time. Right: Velocity of propagation calculated as mm/s traveled by the center of mass of activity (> 2SD) for baseline and stimulation conditions. **C.** Number of repetitions on a next cycle basis per 100 cycles of any propagation pattern for baseline (left) and stimulation (right) conditions across experiments. Note that during baseline a given pattern did not repeat in succession more than four times but repeated up to 40 times in the stimulation condition. Grey area indicated pattern repetition expected by chance. Note that a pattern occurring in larger numbers than a doublet never occurred in our randomized dataset.

Another important change during field application was the repetition of the evoked propagation patterns. During spontaneous conditions, any given propagation pattern could repeat in doublets above what is expected from a randomized dataset and up to a maximum of 4 cycles in a row (Figure 4C – left). However, during field stimulation, activity was much less dynamic with patterns more likely to appear in long sequences; up to 40 consecutive cycles (Figure 4C – right).

## Discussion

In the present study, we have demonstrated that the application of slow sinusoidal alternating electrical fields over frontal regions in the urethane-anesthetized mouse effectively entrained cortical activity and altered global slow-wave dynamics. Under baseline conditions, slow-wave propagation was dynamic with two major (and opposing) patterns along an anterior-lateral to posterior-medial axis. During delivery of alternating electrical fields to frontal regions, slow-wave dynamics showed suppression of the endogenous rhythm in favor of entrainment to a novel slow-wave frequency, corresponding to the frequency of stimulation, that propagated in a novel pattern that showed marked stereotypy. Given that electrical field stimulation has been used to alter neuro-cognitive functions in both online and offline paradigms, our results highlight important mechanistic considerations for how this manipulation might alter spontaneous and ongoing dynamics in a purposed fashion for targeting and altering forebrain interactions.

### Entrainment of cortical activity with sinusoidal electrical field stimulation

We observed entrainment of VSD activity to the stimulation sine-wave which was applied at a faster frequency (1.67 Hz) than the endogenous SO (~ 0.5 – 0.8 Hz). Activity was entrained across the entire cortex. This is in line with previous work from our group and others showing that field stimulation entrains electrical activity at cortical and hippocampal sites, including local-field and unit activity ^10, 33, 34, 35^. Cortical SO activity has also been successfully entrained using transcranial magnetic fields (TMS) ^40^, optogenetic ^41, 42^ and auditory ^11, 43^ stimulation. We also found that during stimulation there was a tendency for VSD spectral and RMS activity to increase at the exact stimulation frequency, corresponding to a decrease in spectral power and RMS level for the endogenous SO frequency. We have previously shown that during field stimulation cortical power and cortico-cortical coherence at the endogenous SO frequency decreases compared to baseline conditions ^33^. This suggests that the applied faster rhythm recruits cortical networks away from their natural participation in the SO rhythm, perhaps creating competition between the externally applied and internally generated rhythms. These results constitute the first evaluation of large-scale activity dynamics during electrical field stimulation, made possible by the negligible levels of artefact generated in VSD recordings which would otherwise swamp typical electrophysiological methods.

### Modulation of large-scale slow-wave cortical propagation dynamics with sinusoidal alternating electrical field application

By tracking and characterizing patterns describing the propagation of slow-wave activity across the surface of the cortex, we showed that the endogenous dynamics of the SO were dramatically altered by stimulation. During baseline spontaneous conditions, we showed a total of ~14 pattern groups with two main and opposing patterns along an anterior-lateral to posterior-medial axis being the most common. Of these two patterns, the anterior-to-posterior direction (AL-PM) was more prevalent in the majority of experiments (9/14) which is consistent with previous research showing mainly anterior-to-posterior SO propagation ^23, 26, 44^. Our observation of multiple possible initiation and termination zones is also highly consistent with previous work in human ^23, 26^ and animal studies ^24, 45, 46, 47^. The initiation and termination zones were highly concentrated in somatosensory as well as visual and retrosplenial areas, which seem to follow major cortical anatomical projections. It has been previously demonstrated that both evoked and spontaneous cortical activity travels along major anatomical routes^30^. The high involvement of the retrosplenial area fits well with the implications of SO in memory consolidation as this region is involved in many high-order cognitive functions including spatial navigation and memory ^48^.

The finding of opposing directional patterns matches well with our previous work using a simple electrode array in rat motor cortex in which we showed that patterns of opposing direction compete for expression ^33^. We also demonstrated here that there is a level of predictability in SO propagation whereby a specific pattern is more likely to repeat on the subsequent cycle, which is also consistent with our previous work ^33^. Beyond the two major patterns, we also observed less common patterns with unique complex features such as multiple initiation and termination zones on a given cycle, propagation constrained to posterior regions and asymmetrical propagation (Supplementary Figure 4). These findings as well are consistent with previous work using both large and fine spatial scale methods ^24, 29, 49^.

The velocity of propagation under spontaneous conditions in our mouse model was ~ 18 mm/s which is slower compared to SO propagation reported for rats (~25-140 mm/s) ^33, 47, 50^, and even more dramatically as compared to humans (> 2 m/s) ^23^. This velocity difference across species can be interpreted as an evolutionary scalable property whereby larger brains make use of disproportionally more long-range cortical connections to compensate for increased distance (but unchanging maximal axonal conduction) between cortical regions ^2, 51^. We further acknowledge that we cannot rule out the possibility that the velocity difference may be a result of the methodological differences between electrophysiological and VSD imaging techniques.

Most interesting were the effects of rhythmic electrical field application on cortical slow-wave dynamics. We demonstrated that field stimulation drastically altered propagation, introducing a stereotyped pattern with varied and distributed initiation zones but with a consistent termination zone in the posterior primary somatosensory cortex. The pattern detected during stimulation was not present during baseline conditions, represented a larger proportion of all detected patterns during stimulation, and tended to repeat for many consecutive cycles. To our knowledge, this is the first demonstration of field stimulation influence on slow-wave dynamics across widespread cortical regions. Previously, we have demonstrated that field stimulation biases SO propagation in an anterior-to-posterior direction using a three-electrode array in rat M1 ^33,39^. Our current findings are consistent with a general interpretation of an anterior-to-posterior direction of propagation during stimulation. However, the limited cortical information in our previous work occluded the description of the complex spatial trajectory of such propagation and its comparison with the spontaneous trajectories across widespread functional regions.

### Implications of altered slow-wave dynamics for memory consolidation

Given that electrical field stimulation during SWS has been shown to improve memory consolidation ^10, 13^, an understanding of how this manipulation might alter slow-wave propagation dynamics is mechanistically relevant for memory processes. During spontaneous SO conditions, we showed a dynamic SO with consistently alternating propagation patterns cycle-to-cycle, while during stimulation, propagation was altered in both its pattern trajectory and its stereotypy. These results may suggest that while spontaneous SO is well suited to consolidate a variety of memories reflected in the activation of many various network interactions from cycle-to-cycle, perhaps the application of field stimulation is more selective for specific (perhaps recently encoded) motifs and subsequently strengthens their relevant functional associations and underlying memories.

In line with this idea, it has been suggested that on every wave the cortical SO may select specific hippocampal cell assemblies, and associated experiences, that will be replayed across hippocampo-neocortical networks ^21, 25, 52, 53, 54, 55, 56, 57^. Field stimulation may be suited to bias these interactions since it has been shown by us and others ^35^ to successfully entrain hippocampal activity likely by way of direct entorhinal input from the temporoammonic path ^34^. We have recently demonstrated that field stimulation increases hippocampal sharp-wave ripple events and cortical spindles and enhances cortico-hippocampal synchrony for both slow and gamma range networks ^34^. Therefore, altering cortical SO propagation using field stimulation likely selects which cortico-hippocampal assemblies are strengthened through interplay. Future work may determine how various stimulation locations may elicit distinct propagation patterns and how those patterns may relate to the targeting of specific memories.

## Conclusion

We demonstrate here for the first time the ability of sinusoidal electrical field stimulation to entrain cortical VSD networks and alter slow-wave propagation dynamics both during and following such manipulation. These results bring us closer to understanding how such stimulation can be used in a targeted manner to enhance or disrupt hippocampal-dependent memory consolidation during offline states.

## Materials and Methods

### Animals

Spontaneous VSD recordings were conducted in 14 adult mice (M = 1, F = 13; C57BL6J = 11, negative Thy1-YFP transgenic strain = 3) with a mean (± standard error of the mean; SEM) weight of 22.8 ± 0.51 g. Of these animals, 8 were used for delivery of sinusoidal field stimulation and for recordings following the cessation of stimulation (≥ 8 min post-stim). Of these 8 animals, 3 were used for assessing post-stimulation effects on a longer timescale (up to 75 min). Multiple and different modality sensory evoked responses and spontaneous recordings were obtained for all animals.

Animals were housed on a 12-hour light/dark cycle and fed a standard mouse diet *ad libitum.* All housing and surgical procedures conformed to the guidelines of the Canadian Council on Animal Care (CCAC) and were approved by the Animal Welfare Committee of the University of Lethbridge.

### Surgery

Mice were initially anesthetized in a gas chamber with a 3% concentration of isoflurane in 100% O_2_ and injected (i.p.) with an initial dose of urethane (1.25 g/kg) and HEPES-buffered saline (100 µL/g). Animals were then transferred to a stereotaxic apparatus and maintained at 0.5-1% isoflurane administered via nose cone. The body temperature was maintained at 37 °C throughout the experiment via a servo-controlled heating pad placed under the mouse. For field stimulation, 4 Teflon-coated stainless steel wires (277 μm bare diameter; 0.4 Ω resistance) were lowered to pial surface and implanted in a triangular configuration with the two central poles located just offset from the midline (+3.5 mm AP and ±0.3 mm ML) and the two lateral poles more posterior and lateral (+2.5 mm AP; ±2.5 mm ML) (see Supplemental Figure 1A). The electrodes were secured to the skull using dental acrylic and super glue.

Mice were then transferred to a surgical plate that was later mounted under the VSD imaging system. A steel head plate (8 X 8 mm) was then mounted onto the skull and a craniotomy window was created by removing the skull (Supplemental Figure 1A). The dura was either removed (n = 7) or kept intact (n = 7). A tracheotomy was performed to aid breathing. Isoflurane delivery was then terminated and additional top-up doses of urethane (10% of initial dose; 0.125 g/kg) were administered every few hours as necessary to maintain a consistent level of anesthesia.

### VSD Imaging

Following craniotomy, the exposed brain surface was bathed in the dye RH1691 (Optical Imaging, New York, NY), which was dissolved in HEPES-buffered saline solution (0.5-1 mg ml^−1^). The staining procedure resulted in a dark purple appearance of the cortex which indicated sufficient dye incorporation as reported previously ^24^. The staining time required for reaching proper incorporation was 30 min – 1 hour for the animals in which the dura was removed and 2.5 – 3 hours for the animals with an intact dura. Following washout of the unbound dye, animals were transferred to the VSD recording rig. The exposed cortex was coated with 1.5% agarose made in HEPES-buffered saline and covered with a glass coverslip to minimize recording artifacts associated with respiration and heartbeat.

VSD images were collected with a CCD camera (1M60 Pantera, Dalsa, Waterloo, ON) and EPIX E8 frame grabber with XCAP 3.8 imaging software (EPIX, Inc., Buffalo Grove IL). The camera was focused ~1 mm dorsoventral to cortical surface to avoid distortion from blood vessels. VSD was excited with a red LED (Luxeon K2, 627 nm centre) and excitation filters (630±15 nm). Images were taken through a macroscope composed of front-to-front video lenses (8.6 X 8.6 mm field of view, 67 µm per pixel). VSD fluorescence was filtered using a 673–703-nm bandpass optical filter (Semrock, New York, NY). Images were acquired in 12-bit resolution at 150 Hz sampling rate for sensory-evoked data and 100 Hz or 200 Hz for spontaneous recordings. The camera-clock output of the EPIX system was recorded for offline alignment of EEG and imaging data.

VSD images were preprocessed using custom-written code in Matlab (Mathworks, Natick, MA). The VSD images were first subjected to principal-component analysis (PCA) and the leading components were retained (15-50 components). The data were then filtered using a second-order Chebyshev bandpass filter (zero-phase filter; 0.2 – 50 Hz). VSD responses were expressed as percent change relative to the baseline response calculated as: F-F0/F0 x 100 where F represents the fluorescence signal for each pixel for a given frame and F0 represents the mean response in each pixel across all frames.

### EEG and ECG recordings

For all experiments a cortical EEG electrode was inserted into the conductive agarose close to the craniotomy window on the right hemisphere near the visual cortex (Supplemental Figure 1A). EEG recordings were stable throughout the experiment and were well correlated with the VSD signal as shown previously ^24^ (Supplemental Figure 2). An electrocardiogram (ECG) electrode was inserted subcutaneously into the back. An electrode inserted on the nasal bone served as a reference for both EEG and ECG signals. All electrodes were teflon-coated silver wires (0.127 mm bare diameter). Signals were wide-bandpass filtered between 0.1 Hz and 1 kHz and amplified at a gain of 1000 using a differential AC amplifier (P5 series Grass, Natus Neurology Inc., Warwick, RI). Signals were digitized using an A-D board (Digidata 1550; Axon Instruments/Molecular Devices, Sunnyvale, CA) and sampled at 2000 Hz to prevent aliasing. Data were acquired and stored using Clampex (Axon Instruments/Molecular Devices) on a PC system for offline analysis.

### Sensory stimulation

Sensory-evoked potentials were collected for all animals to map the cortex. For primary hind limb (HLS1) and fore limb (FLS1) evoked-potentials, brief current pulses (1-2mA, 1 ms; 10 s inter-stimulus interval) were delivered through acupuncture needles (0.14 mm) inserted into all 4 paws. For visual evoked potentials (V1), LED light stimulation (1 ms, green and blue light) was delivered to each eye separately. For barrel cortex stimulation (C2) a single whisker was identified and placed under a piezoelectric device (Q220-A4-203YB, Piezo Systems, Inc., Woburn, MA) that delivered a single tap using a square pulse (1 ms) which moved the whisker 90 μm in an anterior-to-posterior direction. For each type of sensory stimulation, an average of 10 trials was used to locate the position of the primary region. Based on mapping of cortical regions in each animal, VSD data could be represented on a standardized mouse cortical map adapted from the Allen Institute for Brain Science (Seattle, WA) (c.f. Figure 3B).

### Sinusoidal field stimulation

Sinusoidal stimulation was applied using an analogue function generator (Hewlett Packard; 3310A) to the 4 stimulation poles with the two lateral poles receiving opposite polarity stimulation to the two central poles (see Supplemental Figure 1A; ^33, 34^); adapted from ^35^). The stimulation generator output was also digitized using the same system as that used for EEG and ECG signals to obtain stimulation phase and voltage intensity information for later offline analysis. The intensity of stimulation was 10 V (defined as peak-to-peak amplitude). The frequency of stimulation was 1.67 Hz, slightly faster than the endogenous SO frequency (~ 0.5-0.8 Hz). We chose this faster frequency in order to spectrally separate the effects of stimulation and endogenous SO as well as in an attempt to successfully entrain networks by providing a faster putative pace-making stimulus. Stimulation was applied during SO states identified visually in the online EEG recording and was applied for 3-4 min consecutively for one trial. VSD and EEG recordings were obtained before, during, and following stimulation.

In order to measure the amount of current delivered to each hemisphere, we connected a resistor (100 Ω) in series with the stimulation. Measurement of the voltage drop across the resistor allowed us to calculate the current passing through the circuit.

### Pharmacology

In order to ensure that field stimulation did not contaminate the VSD signals, we conducted control experiments in which field stimulation was applied under the influence of three activity suppressing/abolishing conditions: the application of 2% lidocaine (Na^+^ channel blocker, n = 4), combined application of 2mM CNXQ (AMPA receptor antagonist) and 0.1mM MK-801 (NMDA receptor antagonist) (n = 3), as well as during post-mortem recordings following brain death (n = 5). All drugs were dissolved in HEPES-buffered saline solution. For CNQX, a stock solution was created by dissolving CNQX in DMSO, with a final DMSO concentration of 4% in the experimental solution. After collection of baseline and field stimulation recordings, the agarose was removed from the surface of the brain (for the dura-intact animals) and lidocaine (2%) was applied for ~30 min (until suppression was evident in the EEG) before the trial began. Lidocaine was then washed out for ~60 min (until resumption of normal EEG activity) and then the mixture of CNQX (2 mM) and MK-801 (0.1 mM) was applied for ~45 min (until suppression was evident in the EEG). Following the CNQX+MK-801 condition animals were euthanized with euthasol (0.05 – 0.1 mL) and the final control trial began immediately after flattening of both the EEG and ECG signals. For each of these activity suppressing conditions the trial began with 3-4 min of spontaneous VSD and EEG recording followed by 3-4 min of electric field stimulation. For all experimental conditions imaging took place while the cortex was bathed in the drug-containing buffer to maintain consistent suppression effects. Agarose was not reapplied. Heartbeat artifacts were thus present during drug trials at ~ 8 Hz but did not affect our frequency of interest containing the slow oscillation and stimulation frequencies (~0.5-1.67 Hz) (Supplemental Figure 2C).

### Data analysis

All analyses were performed using a combination of built-in functions and custom written code in Matlab (MathWorks). All means are reported as mean ± standard error of the mean (SEM). Significance for all statistical tests was defined as p < 0.05 and all *t* tests were performed two-tailed and between-subjects (due to uneven subject numbers per condition) unless otherwise specified.

Spectral analyses were performed using Welch’s averaged modified periodogram method for power spectral density estimation (6 s window; 2 s overlap) calculated on continuous 120 second epochs of data that were then averaged based on subject and condition. For VSD data, this method was performed on data taken from a given region of interest (ROI) after averaging activity from 25 pixels (5 x 5 window) (Supplemental Figure 1C).

The root-mean square (RMS) level was computed for each pixel of the VSD data over a 3-minute period for each condition within each experiment. To create group averages, RMS values were first averaged across all pixels for either one (Supplementary Figure 8) or both (Supplemental Figure 2) hemispheres within each experiment and condition.

For assessing the level of VSD activity entrainment to the delivered stimulus sine-wave, stimulus-triggered VSD averages were created. The phase angle of the Hilbert transform yields a measure of instantaneous phase over time and was used to identify trigger points (Figure 1B) ^33^. To obtain a baseline measure of VSD activity entrainment to the endogenous slow oscillation, the positive peak of the slow oscillation (90º) at a given ROI was used to trigger VSD traces across all cycles (Supplementary Figure 7A – left). For the field stimulation conditions, the positive peak of the delivered sine-wave was used as the trigger point. The peak-to-trough amplitude (in SD units) was taken as a measure of entrainment for all conditions.

### Propagation patterns

In order to identify propagation patterns on a cycle-by-cycle basis, we first filtered VSD activity for every pixel around the spectral peak of activity (0.3 – 1.1 Hz for baseline, 1.1 – 1.9 Hz for stimulation). The phase angle of the Hilbert transform was used to identify positive peaks in the VSD signal taken at the ROI of HLS1 (identifying cycles from the EEG SO instead yielded identical results) or taken from the positive peaks of the stimulation sine wave for stimulation conditions (Figure 1A)^33^. Using this trigger point, VSD images were separated into single activation (ON state) cycle periods defined as ±135º from the positive peak. The phase angle of the Hilbert transform was then computed for every pixel within each cycle and the positive peak of the given pixel was identified. Thus, for every cycle, a “propagation” image could be created which shows the time (ms) at which each pixel reached its positive peak (peak activation) relative to the triggered peak taken at the ROI (time = 0). By assessing the earliest and latest peak onsets in this image a propagation pattern could be inferred (c.f. Figure 1D – gray images). These single cycle propagation images were then subjected to a hierarchical clustering algorithm (built-in Matlab toolbox) in order to group patterns with similar spatial profiles. To create a similarity measure used for grouping, each pair of patterns was spatially correlated and these correlation measures were used in the algorithm. A pattern group was defined as one in which any 2 patterns within the group had a spatial correlation of > 0.08 (above the 99^th^ percentile from 1000 trials of randomized data) resulting in the identification of ~14 pattern groups during baseline (Figure 4A). Patterns across all conditions were subjected simultaneously to the clustering algorithm in order to identify overlapping pattern groups across conditions if such existed.

Start and end points of propagation for each cycle were defined as the center of mass of the pixels with the earliest and latest Hilbert positive peak detections (< 2SD and > 2SD respectively) and directional vectors could be drawn from start to end point for those patterns with a relatively linear propagation direction. A measure of propagation strength (propagation index) was calculated as the change in area of activation (> 2SD above baseline fluorescence) from frame to frame (Δmm^2^/frame). The velocity of propagation for any given cycle was calculated as total distance traveled from start to end point over time (mm/s).

## Code Availability

All custom written code in MATLAB used in this work is available upon request.

## Data Availability

All data collected and used in this work is available upon request.

## Acknowledgments

This work was supported by Natural Science and Engineering Council of Canada (NSERC) grants #249861 to CTD and #40352 to MHM. MHM is a Campus Alberta for Innovation Program Chair, Alberta Alzheimer Research Program (MHM), and Alzheimer Society of Canada (MHM). AG was supported by an NSERC Alexander Graham Bell Canada Doctoral Graduate Scholarship and a grant from the Campus Alberta Neuroscience Trainee Mobility Program. JKA was supported via an NSERC CREATE (Biological Information Processing) Scholarship. We thank Dr. Jianjun Sun for surgical assistance.

